# A novel prokaryotic CRISPR-Cas12a based tool for programmable transcriptional activation and repression

**DOI:** 10.1101/2020.08.05.232744

**Authors:** Christoph Schilling, Mattheos A.G. Koffas, Volker Sieber, Jochen Schmid

**Author notes:** Abbreviations: CRISPR: clustered regularly interspaced short palindromic repeats; CRISPRi: CRISPR interference; CRISPRa: CRISPR activation; gRNA: guide RNA; GTi: initiating glycosyltransferase; 2,3-BDL: 2,3-butanediol; EPS: exopolysaccharide; *bdh*: butanediol dehydrogenase; *ldh*: lactate dehydrogenase.

## Abstract

Transcriptional perturbation using inactivated CRISPR-nucleases (dCas) is a common method in eukaryotic organisms. While rare examples of dCas9 based tools for prokaryotes have been described, multiplexing approaches are limited due to the used effector nuclease. For the first time, a dCas12a derived tool for the targeted activation and repression of genes was developed. Therefore, a previously described SoxS activator domain was linked to dCas12a to enable programmable activation of gene expression. Proof of principle of transcriptional regulation was demonstrated based on fluorescence reporter assays using the alternative host organism *Paenibacillus polymyxa* as well as *Escherichia coli*. Single target and multiplex CRISPR interference targeting the exopolysaccharide biosynthesis of *P. polymyxa* was shown to emulate polymer compositions of gene knock-outs. Simultaneous expression of 11 gRNAs targeting multiple lactate dehydrogenases and a butanediol dehydrogenase resulted in decreased lactate formation, as well as an increased butanediol production in microaerobic fermentation processes. Even though Cas12a is more restricted in terms of its genomic target sequences compared to Cas9, its ability to efficiently process its own guide RNAs *in vivo* makes it a promising tool to orchestrate sophisticated genetic reprogramming of bacterial cells or to screen for engineering targets in the genome. The developed tool will accelerate metabolic engineering efforts in the alternative host organism *P. polymyxa* and might be also applied for other bacterial cell factories.

---

Seeking a biobased and sustainable economy, bacterial cell factories have been used for the production of a variety of high value products such as amino acids^1^, biofuels^2^ or biosynthesis of complex pharmaceutical compounds like artemisinic acid^3^. The development of robust production strains for industrial scale production typically requires a deep understanding of the underlying metabolic networks enabling sophisticated engineering technologies to optimize fluxes towards the product of interest and eliminating unwanted side products^4^. Within the last decade, the development of new technologies such as CRISPR-Cas9 mediated genome editing resulted in a dramatic increase in the complexity and scope of metabolic engineering approaches^5–8^.

CRISPR interference (CRISPRi), using catalytically inactive CRISPR nucleases (dCas) for transcriptional repression, has been successfully demonstrated in bacteria^9–11^. In eukaryotic organisms, CRISPRi has been further expanded by direct fusion of Cas9 and effector domains to remodel the chromatin structure of target genes resulting in an even tighter repression of expression^12–14^. Contrary, CRISPR-mediated activation (CRISPRa) of gene expression was achieved by linking the deactivated CRISPR-nuclease from *Streptococcus pyogenes* dCas9 to transcriptional activation domains via translational fusion or recruitment via peptide epitopes or additional RNA scaffolds^15–17^. While CRISPRa has been extensively used and further developed for eukaryotic organisms to activate transcription of target genes^18^, the number of synthetic tools for prokaryotes is still limited. Recently, new CRISPR-Cas9 based systems were developed for bacteria using effector domains such as RpoZ^9^, RpoA^19,20^, bacteriophage derived transcriptional activators like AsiA^21,22^ or the more effective AraC family transcription factor SoxS^23^ that facilitate the recruitment of the RNA polymerase holoenzyme. In order to overcome narrow target site requirements, more flexible CRISPRa toolkits using σ^54^-dependent promoters were established^24^. While σ^70^-activators bind in close proximity to its cognate promoter and recruit RNA-polymerase, bacterial enhancer binding proteins (bEBPs) corresponding to σ ^54^-dependent promoters target long distance upstream activating sequences (UAS) and initiate transcription similar to eukaryotic promoters^24^. Consequently, CRISPR-dCas guided bEBPs were directed to UAS regions in order to enable a more flexible, long distance regulation of target promoters resulting in a remarkable dynamic output range ^24^.

While Cas9 derived from *Streptococcus pyogenes* is the most well-studied RNA-guided endonuclease and was used in a multitude of studies, it has demonstrated several downsides in simultaneously targeting multiple loci. Although, multiplex genome editing can be realized, a uniform expression of multiple gRNAs proved to be challenging. Strategies to overcome this constraint include the expression of sgRNA transcripts from multiple plasmids, the co-expression of RNA processing enzymes such as RNAse III^25^ and Csy4^26,27^ or flanking of consecutive gRNAs by ribozymes or tRNAs that enable efficient processing of the mature gRNA^28,29^ from a single transcript. However, all these strategies are limited in the number of multiplex targets due to cytotoxic effects. Contrary, multiplex genome editing approaches using Cas12a nuclease orthologs (also known as Cpf1) from *Francisella novicida, Acidaminococcus* sp. or *Lachnospiraceae* sp. require only the expression of a single crRNA array^30,31^., Unlike Cas9, Cas12a additionally possesses RNase activities to process the precursor crRNA array and form the gRNAs necessary to direct the CRISPR nuclease to the target DNA^31^. Leveraging this dual RNase/DNase function, simultaneous perturbation of 25 individual targets was demonstrated in mammalian cell lines using a single transcript harboring both, the open reading frame of Cas12a and a CRISPR array^32^. In contrast to Cas9, the PAM is located upstream of the cleavage site and consists of a sequence with a very low GC content. For all commonly used Cas12a nucleases from *Francisella novicida, Acidaminococcus* sp. or *Lachnospiraceae* sp. the most efficient protospacer adjacent motif (PAM) required for cleavage was determined as TTTV^33,34^. While this particular PAM is more restrictive compared to NGG of *Sp*Cas9, protein engineering efforts to loosen the stringency of CRISPR nucleases to enable genome editing in otherwise inaccessible loci were successful^35,36^.

Due to this advantages for multiplex genome perturbation studies, dCas12a has been extensively used for the tunable transcriptional regulation of gene expression via CRISPRi and CRISPRa in eukaryotic cells^30,32,36^. Despite rapid advances in CRISPR-based technologies, to the best of our knowledge, only CRISPR interference studies have previously been reported for prokaryotic cells using dCas12a^37^, while publications demonstrating the targeted gene activation via CRISPRa are still missing for this promising approach. Therefore, the aim of this study was to establish a functional dCas12a based multiplex gene modulation system capable of CRISPRa and CRISPRi using a broad-host range plasmid.

*Paenibacillus polymyxa* is a Gram-positive, spore forming, non-pathogenic, soil bacterium^38^ of biotechnological interest for its ability to produce enantiopure *R,R*-2,3-butanediol (2,3-BDL) and exopolysaccharides (EPS) with interesting material properties^39,40^ (Figure S1). *P. polymyxa* DSM 365 putatively produces two distinct heteroexopolysaccharides we termed Paenan I and Paenan II, which is reflected in two functionally complete EPS clusters encoding all genes required for the Wzx/Wzy biosynthesis pathway^41,42^. Knock-outs of distinct glycosyltransferases within the clusters resulted in EPS variants with altered rheological behavior^41^. *P. polymyxa* is also applied in the production of 2,3-BDL via the mixed-acid fermentation pathway in microaerobic conditions. Depending on oxygen availability, production of side products such as lactate, formate and ethanol is required to maintain the redox balance^43^. We recently showed that targeted engineering of the side pathways competing for pyruvate significantly increased the productivity of 2,3-BDL biosynthesis^44^. However, the knock-out of a specific lactate dehydrogenase, did not result in decreased titers of the by-product lactate due to the action of redundant homologs. In this study, lactate should be eliminated via the concerted knock-down of all four different lactate dehydrogenases found in the genome of *P. polymyxa*. Additionally, the carbon flux should be directed towards 2,3-BDL by inducing the expression of butanediol dehydrogenase (*bdh*) in parallel via a newly developed CRISPRa/i system.

## Materials and Methods

### Strains and media

*P. polymyxa* DSM 365 was acquired from the German Collection of Microorganisms and Cell Culture (DSMZ, Germany). *E. coli* NEB Turbo cells (New England Biolabs, USA) were used for any plasmid construction presented in this study. *E. coli* S17-1 (DSMZ strain DSM 9079) was utilized for transformation of *P. polymyxa* DSM 365 via conjugation. All medium components were obtained from Carl Roth GmbH (Germany) if not indicated differently. For cloning procedures, strains were grown in LB media (5 g L^-1^ yeast extract, 10 g L^-1^ tryptone, 10 g L^-1^ NaCl) and additionally supplemented with 50 µg mL^-1^ neomycin and 20 µg mL^-1^ polymyxin if required. All strains were stored in 30 % glycerol at −80 °C. Prior to cultivation, strains were streaked on LB agar plates and grown at 30 °C. All strains used or constructed in this study are listed in Table S1. For 2,3-BDL fermentations a single colony was used for inoculation of 50 mL pre-culture medium containing 60 g L^-1^ glucose, 5 g L^-1^ yeast extract, 5 g L^-1^ tryptone, 0.2 g L^-1^ MgSO_4_ heptahydrate (Sigma Aldrich, USA), 3.5 g L^-1^ KH_2_PO_4_, 2.5 g L^-1^ K_2_HPO_4_. Fermentation medium components were autoclaved separately and contained 120 g L^-1^ glucose, 5 g L^-1^ yeast extract, 3.5 g L^-1^ tryptone, 0.2 g L^-1^ MgSO_4_x7 H_2_O, 3.5 g L^-1^ KH_2_PO4, 2.5 g L^-1^ K_2_HPO_4_, 5 g L^-1^ ammonium acetate, 4 g L^-1^ (NH_4_)_2_SO_4_ and 3 mL L^-1^ trace element solution. Trace element solution contained 2.5 g L^-1^ iron sulfate heptahydrate, 2.1 g L^−1^ sodium tartrate dihydrate, 1.8 g L^−1^ manganese chloride dihydrate, 0.075 g L^−1^ cobalt chloride hexahydrate, 0.031 g L^−1^ copper sulfate pentahydrate, 0.258 g L^−1^ boric acid, 0.023 g L^−1^ sodium molybdate dihydrate and 0.021 g L^−1^ zinc chloride. Trace element solution was filter-sterilized and added to the media after cooling down to room temperature.

For EPS production, MM1 P100 medium^39^ was used as described before, containing 30 g L^-1^ glucose and 5 g L^-1^ peptone. The corresponding pre-culture medium contained a reduced amount of 10 g L^-1^ glucose and was buffered to pH 6.8 with 20 g L^-1^ MOPS.

### Plasmid construction

The gene encoding for an engineered (E174R, N282A, S542R, K548R)^36^ catalytically inactivate (D908A) variant of AsCas12a was codon optimized for *Bacillus* ssp. and synthesized by ATG:biosynthetics (Germany). The basic plasmid pCRai (Figure S2) was assembled by isothermal Gibson Assembly^45^ from three PCR-amplified fragments consisting of a pUB110 derived backbone including oriT for conjugational transfer^41^, a lacZ replacement cassette for BbsI based cloning of target gRNAs and the codon optimized *enAsdcas12a* cassette. Activator domains were PCR-amplified from extracted gDNA of *P. polymyxa* DSM 365 and *E. coli* NEB Turbo respectively and cloned into pCRai by Golden Gate Assembly using BsaI. Cloning of gRNA sequences was conducted as previously described^41^. The PsgsE-sfGFP reporter was cloned via isothermal Gibson Assembly using a unique SpeI site of pCRai. The dual reporter plasmid was constructed by cloning a PsgsE-mRFP and a PsgsE-sfGFP reporter cassette in tandem by Golden Gate Assembly after linearization of pCRai_soxS with SpeI/SalI. All oligonucleotides used for the construction of plasmids are listed in Table S2 and S3. gRNA targeting sequences (spacers) and DNA sequences of used activator domains are listed in Table S4 and S5.

### Conjugation based transformation of *P. polymyxa* DSM 365

*P. polymyxa* was transformed by conjugation using *E. coli* S17-1 harboring the various plasmids. Overnight cultures of donor and recipient strains were diluted 1:100 with selective or non-selective LB media respectively and cultivated at 37 °C for 3 h, 280 rpm. 900 µL of the recipient culture was heat shocked at 42 °C for 15 min and mixed with 300 µL of the donor strain culture. Cells were centrifuged at 6,000 g for 2 min, resuspended in 800 µL LB media and dropped on non-selective LB agar plates. After 24 h of incubation at 30 °C, cells were scrapped off, resuspended in 500 µL LB-broth and 100 µL thereof plated on selective LB-agar containing 50 µg mL^-1^ neomycin and 20 µg mL^-1^ polymyxin for counter selection. *P. polymyxa* conjugants were analyzed for successful transformation after 48 h incubation at 30 °C by cPCR. Confirmed knock out strains were plasmid cured by cultivation in LB broth without antibiotic selection pressure and subsequent replica plating on LB agar plates both with and without neomycin. Strains that did not grow on plates with selection marker were verified by sequencing of the target region and used for further experiments.

### Photometric assay

For sfGFP fluorescence experiments, 3 mL of EPS medium supplemented with 50 µg mL^-1^ neomycin were inoculated with a single colony of the respective strains and grown over night at 30 °C, 200 rpm. After 18 h, each strain was sub-cultured 1:100 in 3 mL selective MM1 P100 medium and grown for 24 h at 30 °C, 200 rpm. After 24 h, 100 µL were transferred to a 96 well microtiter plate and OD_600_, GFP fluorescence (Ex. 488 nm Em. 515 nm) and mRFP fluorescence (Ex. 560 nm Em. 600 nm) measured in a Ultraspec 10 spectrophotometer (Amersham Biosciences, UK). Fluorescence values were normalized to OD_600_ in all experiments. In parallel, 1 mL of each culture was pelleted by centrifugation and used for qPCR experiments. Relative expression signals were calculated based on an off-target construct expressing gRNAs not binding to the genome of *P. polymyxa* DSM 365 or any plasmid used in this study (Table S 4).

### Quantitative RT-PCR

RNA extraction of positive samples of the GFP fluorescence assay as well as butanediol fermentation processes was performed using the Aurum Total RNA Mini Kit (BioRad, USA) according to the manufacturer’s instructions. Synthesis of cDNA was conducted using iScript reverse transcriptase (BioRad, USA) using 1 µg total RNA template. The qPCR reactions were performed in triplicates on a CFX-96 thermocycler using SsoAdvanced Universal SYBR Green Supermix (BioRad, USA) using 5 ng of cDNA as a template in 10 µL reaction volume. Negative controls without reverse transcriptase during cDNA synthesis were used in order to evaluate the absence of gDNA contaminations. Relative gene expression levels were calculated based on the ΔΔCq method^46^ and *gyrA* as a reference gene. After qPCR, a melting curve analysis was performed to confirm the presence of a single PCR product for each target. Designed primers were analyzed by the OligoAnalyzer Tool (IDT, USA) to avoid hairpin formation, self- and hetero dimer formation with free energy values more than 10 kcal mol^-1^. Oligonucleotides used for qPCR experiments are listed in Table S2.

### CRISPR-Cas9 mediated genome editing

All gene knock-outs were performed as previously described by Rütering et al.^41^. In brief, gRNAs for the targeted genome regions were designed using Benchling CRISPR Design Tool. For each target a minimum of two gRNAs were designed typically targeting distinct regions of the open reading frame. Oligonucleotides were phosphorylated, annealed and cloned into pCasPP by Golden Gate assembly. Approximately 1 kB up- and downstream homology flanks for each targeted nucleotide sequence were amplified from genomic DNA of *P. polymyxa* DSM 365 using Phusion Polymerase according to the manufacturer’s instructions and fused by overlap extension PCR via a 20 bp overlap. Homology flanks were cloned into pCasPP through Gibson Assembly or molecular cloning after linearization by use of SpeI. After transformation of *E. coli* NEB Turbo, clones were analyzed for correct construct assembly by colony PCR (cPCR) and sequencing of the amplicons. Finally, correct constructs were transferred to chemical competent *E. coli* S17-1 cells for the following conjugational transformation of *P. polymyxa*.

### EPS batch fermentation

EPS fermentations were conducted in a 1 L DASGIP parallel bioreactor system with a working volume of 500 mL. A single colony from a freshly streaked plate was used to inoculate 100 mL MM1 P100 pre-culture medium by following incubation for 16 h at 30 °C, 160 rpm. Bioreactors were inoculated to give an initial OD of 0.1. Fermentation was performed at 30°C and stirrer speed (200 - 600 rpm) and gassing (6 - 10 L h^-1^) with pressurized air through a L-sparger were controlled to maintain 30 % DO saturation. The stirrer was equipped with a 6-plate-rushton impeller placed 2.5 cm from the bottom of the shaft. The pH value was maintained at 6.8 and automatically adjusted with 2 M NaOH or 1.35 M H_3_PO_4_ as required. Foam control was performed using 1 % of antifoam B (Merck, Germany). For monitoring the process parameters, reactors were equipped with redox and dissolved oxygen probes.

After the end of the process the fermentation broth was diluted 1:10 with dH_2_O and the biomass was separated by centrifugation (15 000 g, 20°C, 20 min) followed by cross-flow filtration of the supernatant using a 100 kDa filtration cassette (Hydrosart, Sartorius AG, Germany). EPS was precipitated by slowly pouring the concentrated fermentation supernatant into two volumes of isopropanol. EPS was collected and dried overnight in a VDL53 vacuum oven at 40°C (Binder, Germany). Dry weight of the obtained EPS was determined gravimetrically prior to milling to a fine powder in a ball mill at 30 Hz for 1 min (Mixer Mill MM400, Retsch GmbH, Germany).

### Carbohydrate fingerprinting

EPS monosaccharide composition was analysed using the 1-phenyl-3-methyl-5-pyrazolone high-throughput method (HT-PMP) as previously described using 1 g L^-1^ reconstituted EPS solutions^47^. In brief, 0.1 % EPS solutions were hydrolyzed in a 96 well plate, sealed with a rubber mat and further covered by a custom-made metal device with 2 M trifluoroacetic acid (90 min, 121°C). Samples were neutralized with 3.2 % NH_4_OH. 75 µL of PMP master mix (125 mg PMP, 7 mL MeOH, 3,06 mL dH_2_O, 437.5 µL 3.2% NH_4_OH) were mixed with 25 µL of neutralized hydrolysate and incubated at 70°C for 100 min in a PCR cycler. 20 µL of derivatized samples were mixed with 130 µL of a 1:26 dilution of 0.5 M acetic acid and filtered with a 0.2 µm filter plate (1,000 g, 2 min) followed by HPLC-UV-MS analysis as previously described^47^.

### Butanediol batch fermentation

Batch fermentations were conducted in 1 L DASGIP bioreactors (Eppendorf, Germany) with an initial volume of 550 mL. A single colony from a freshly streaked plate was used to inoculate 100 mL pre-culture medium by following incubation for 16 h at 30 °C, 160 rpm. 50 mL of this cultivation broth (diluted with pre-culture medium if required) were used to inoculate the bioreactor by an initial OD_600_ of 0.1. Fermentation was performed at 35 °C and constant aeration of 0.075 vvm. The stirrer was equipped with a 6-plate-rushton impeller placed 4 cm from the bottom of the shaft and constantly stirring at 300 rpm. The pH value was maintained at 6.0 and automatically adjusted with the addition of 2 M NaOH or 1.35 M H_3_PO_4_ as required. Foam control was performed using 1 % of antifoam B (Merck, Germany). In order to monitor process parameters, reactors were equipped with redox and pH probes. Glucose and product concentrations were determined via a HPLC-UV-RID system (Dionex, USA) equipped with Rezex ROA-H^+^ organic acid column (300 mm x 7.8 mm Phenomenex, USA). Column temperature was set to 70 °C and 2.5 mM H_2_SO_4_ was used as the mobile phase with a flow rate of 0.5 ml min^-1^. All measured concentrations of 2,3-BDL in this publication represent solely the levo-stereoisomer of the alcohol if not explicitly noted differently.

## Results and Discussion

### Identification of functional transcriptional activator domains

In a first step, transcription activator domains of different regulatory protein families were tested in order to identify a suitable candidate for CRISPRa (Table S5). Each domain was linked by translational fusion to the C-terminal end of dCas12a through a 10 amino acid flexible linker peptide (-GSEASGSGRA-). As endogenous transcription activators from *P. polymyxa*, the cAMP receptor protein (CRP) RNA polymerase subunits σ^70^ (RpoD) and ω (RpoZ), as well as the regulator of the glutamate synthase operon (GltC) were evaluated. SoxS, an activator of the superoxide stress genes from *E. coli* was chosen as an additional heterologous regulator. Out of these, only RpoZ and SoxS have previously been reported as suitable candidates using dCas9 based CRISPRa systems^9,23^. The plasmid pCRai_sfGFP was constructed by isothermal assembly based on the previously established Cas9 genome editing plasmid pCasPP^41^. pCRai_sfGFP encodes dCas12a linked to the different transcriptional activators, the corresponding gRNA expression cassette, as well as *sfgfp* under the control of constitutive promoters.

The sgsE-promoter from *Geobacillus stearothermophilus* used for sfGFP expression is a temperature sensitive promoter containing three core promoter sites^48^. At low temperatures of 28 °C the front most core promoter (P3) is active, resulting in a weak basal expression, while elevated temperatures of 37 - 45°C lead to highly increased expression from RNA-polymerase binding sites further upstream (P1 and P2) as shown in *B. subtilis*^48^. Furthermore, this well characterized promoter has demonstrated robust but weak expression for all cultivation conditions used in this study^41,48^. Therefore, the sgsE-promoter was chosen to test CRISPRa activities of the different activator domains in order to induce strong expression levels even at low temperature (Figure 1 A). Eukaryotic CRISPRa systems allow a relatively broad range, in which gRNAs mediate the binding of the CRISPR effector module to efficiently activate or repress the expression of target genes. Contrarily, bacterial CRISPRa systems have demonstrated to be highly sensitive to the correct distance of the activated dCas12a target sequence to the promoter^23^. Bacterial CRISPRa systems act by facilitating the recruitment of the RNA-polymerase to the promoter, while eukaryotic systems typically cause chromatin re-arrangements to interfere with the expression of target genes^13^. For bacterial dCas9 based systems, an optimal distance was determined in the range between 60 to 100 bp upstream of the transcriptional start site (TSS)^23^. The optimal distance might vary depending on different activator domains. Consequently, we tested four different spacers (target sequence of the gRNA) allowing the activated effector module to bind to the template and non-template strand in the range of 40 to 120 bp upstream of the TSS to induce expression from the strong RNA-polymerase binding site P1 (Figure 1). PAM sites for all spacers showed the same motif (TTTC). In order to test whether observed effects actually arise from the binding of the dCas12a-activator complex to the respective target sites, additional constructs expressing off-target gRNAs not binding to the plasmid or genome of *P. polymyxa* DSM 365 were constructed.

**Figure 1:**
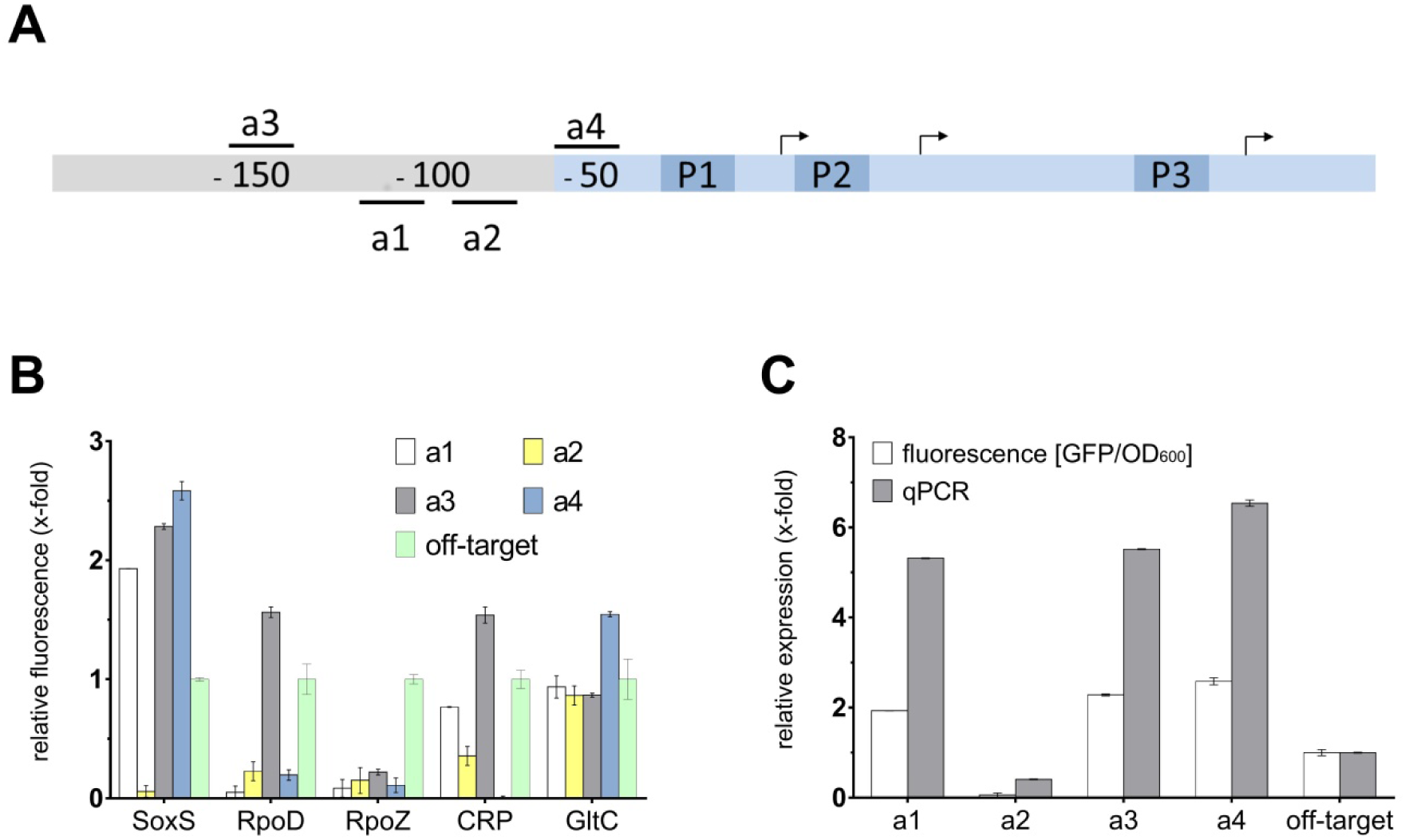
Establishment of a CRISPRa system using dCas12a linked to activator domains. A) Schematic overview of spacers used for CRISPRa (a1-a4) upstream of the *sgsE* promoter (blue). dCas12a was fused to different activator domains (SoxS, RpoD, RpoZ, CRP, GltC) and positioned upstream of the *sgsE*-promoter with multiple spacers on the template and non-template strand. The promoter consists of three core promoter binding sites (P1 - P3), of which the heat-inducible P1 site corresponds to the strongest expression^48^. CRISPRa experiments aim to activate expression from P1 already at low temperatures. Arrows indicate the TSS of the corresponding core promoter sites. B) GFP expression using different activator domains and spacer sequences (a1 - a4) relative to a corresponding off-target gRNA. SoxS showed up to 2.5-fold GFP fluorescence with three gRNAs, while RpoD, CRP and GltC demonstrated elevated fluorescence for only one spacer respectively. C) Expression levels determined by qPCR (relative expression) showed up to 6.5-fold increased transcription levels of *gfp* for soxS variants compared to off-target gRNAs.

Out of all tested activator constructs, *Ec*SoxS demonstrated the best performance in *P. polymyxa* (Figure 1 B). Three out of four tested gRNAs significantly increased expression of sfGFP and showed an increased fluorescence signal during photometric evaluation. The qPCR experiments showed up to 6.5-fold increased transcription levels using gRNA_a4, but also gRNA_a1 and gRNA_a3 bound to dCas12a-soxS displayed increased transcription and fluorescence signals (Figure 1 C). Surprisingly, the highest fluorescence signal was achieved using the spacer a4, which was positioned 50 bp upstream of the TSS of the heat-inducible promoter site P1 (Figure 1 A). While the close proximity of the binding site of this gRNA lies outside of the ideal distance determined for a dCas9-soxS construct^23^, the distance to the second RNA-polymerase binding site (P2) of 85 bp might result in transcription from the secondary heat-inducible promoter.

Additionally, dCas12a fused to other activator domains such as RpoD, GltC and CRP respectively also demonstrated increased fluorescence results for individual spacer sequences (Figure 1 B). However, some combinations also led to a decreased fluorescence signal of GFP in *P. polymyxa* indicating that the effector module blocks the binding of the RNA-polymerase to the promoter and therefore effectively represses transcription of the gene of interest. For GltC and CRP, the use of gRNAs a3 and a4, which are located in close proximity to each other (10 bp), effects on GFP expression changed from 2-fold increased GFP signal to transcriptional repression. All of the investigated activators act by direct interaction with the RNA-polymerase^49–51^. Contrary to other activators, there is experimental evidence suggesting that SoxS already forms a binary pre-recruitment complex with the C-terminal domain of the α-subunit and scans DNA for cognate SoxS binding sites^52,53^. Therefore, we hypothesize that a similar pre-recruitment is formed with the dCas12a-SoxS fusion protein, which allows more flexibility in the correct distancing of the gRNA to the promoter binding site. Due to the consistent performance of the dCas12a-soxS constructs, this particular activator domain was used for all further experiments.

Interestingly, while all experiments using the fluorescence reporter system were performed in *P. polymyxa*, observed effects were almost identical in *E. coli* S17-1 that was used for the conjugational transformation of *P. polymyxa* DSM 365 (Figure S 3). Consequently, we demonstrated a broad-host range use of the constructed pCRai_soxS plasmid in both Gram-positive, as well as Gram-negative bacteria. In case of the fluorescence reporter assays, in which all functional parts were encoded on a single plasmid, it was possible to accelerate the screening of potential guide RNAs by using *E. coli* S17-1 as a pre-screening platform prior to the more time-consuming conjugational transformation of *P. polymyxa* DSM 365.

Our results exemplified that the stringency of gRNA positioning with the SoxS domain is lower compared to other activators. Empirical testing of multiple spacers is still required to enable improved activation of target promoters. However, it might be possible to establish a design rule set to enable *a priori* construction of optimized spacers with more experimental data using different promoters.

### Establishment of CRISPRi and multiplexing CRISPRi/a

In a next step, we evaluated whether the use of the dCas12a-soxS activator constructs is also possible for CRISPRi by re-positioning the activated dCas12a-gRNA complex within the open reading frame of sfGFP. Thereby, the effector module acts as a road block for the RNA-polymerase and inhibits the elongation of the nascent transcript. Three different spacer sequences were tested (Figure 2 A). While expression levels of *sfgfp* were significantly decreased by approximately 80 % for all constructs compared to an off-target construct, the actual fluorescence of GFP remained at higher levels using gRNAs T1 and T2 (Figure 2 A B). Even though, fluorescence signals were not fully eliminated, a severe decrease of up to 95 % using gRNA T3 was observed.

**Figure 2:**
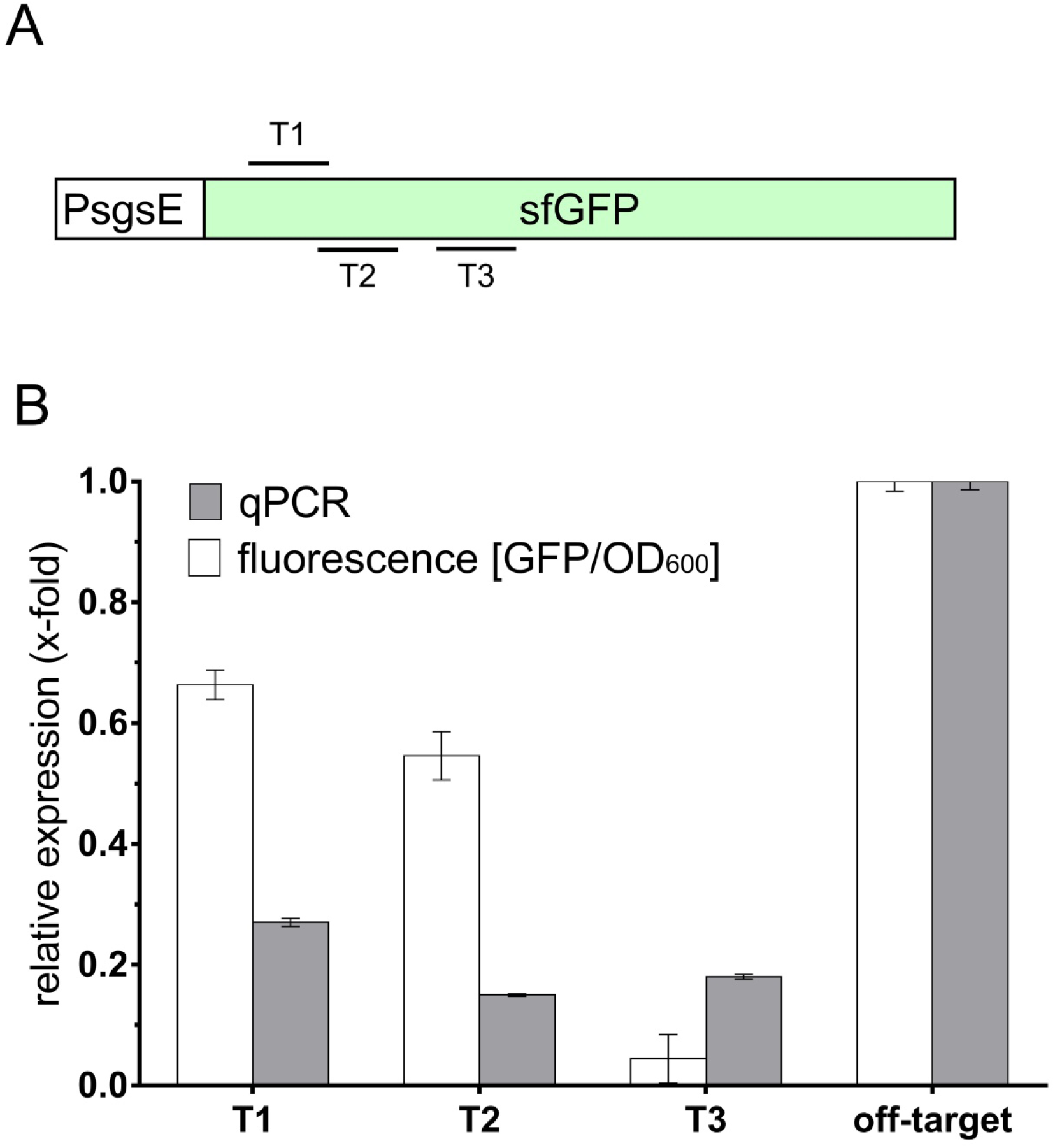
Establishment of CRISPRi in *P. polymyxa* using pCRai_soxS. A) Schematic overview of gRNA binding sites within the ORF of sfGFP. B) Three spacers within the ORF of *sfgfp* (T1-T3) were tested respectively by fluorescence experiments and qPCR expression analysis relative to an off-target gRNA. While transcriptional expression levels were reduced by 75 % to 80 % for all gRNAs, measured fluorescence levels of GFP fluctuated more between different spacers. The activated complex using gRNA T3 demonstrated the best repression resulting in a highly reduced fluorescence signal as well as a reduced transcription of *sfgfp*. Reporter expression was determined photometrically and by qPCR experiments in biological triplicates.

In order to test the capability of our construct to simultaneously repress and activate different target genes, a constitutively expressed mRFP reporter cassette was cloned in pCRai_soxS in addition to the sfGFP reporter. For CRISPRa, the previously used gRNA_a1 was chosen to induce the expression of sfGFP. For CRISPRi a new spacer was designed binding within the ORF of *mrfp* (Figure 3 A). All strains of *P. polymyxa* were compared to a strain harboring an off-target CRISPR-array. Due to weak fluorescence signals after 24 h of inoculation, only transcriptional expression levels were determined via qPCR at this point of time, but photometric evaluation of the reporters was performed after 48 h. When expressed individually, CRISPRi resulted in a reduction of the mRFP fluorescence signal by 74 %, while CRISPRa increased sfGFP expression by 68 % (Figure 3 B). Simultaneous expression of both activated gRNAs from a single CRISPR-array decreased mRFP fluorescence by 60 %, while increasing the fluorescence signal of sfGFP by 120 %. Therefore, we demonstrated the efficient control of the expression of multiple genes using a single CRISPR-array. Depending on the positioning of the spacer, it proved possible to activate or repress multiple target genes in parallel. Interestingly, repression of mRFP alone seemed to positively influence the expression of sfGFP. We hypothesize that this synergistic effect resulted out of a lower competition of the PsgsE promoter corresponding to *sfgfp* and the PsgsE-*mrfp* expression cassette.

**Figure 3:**
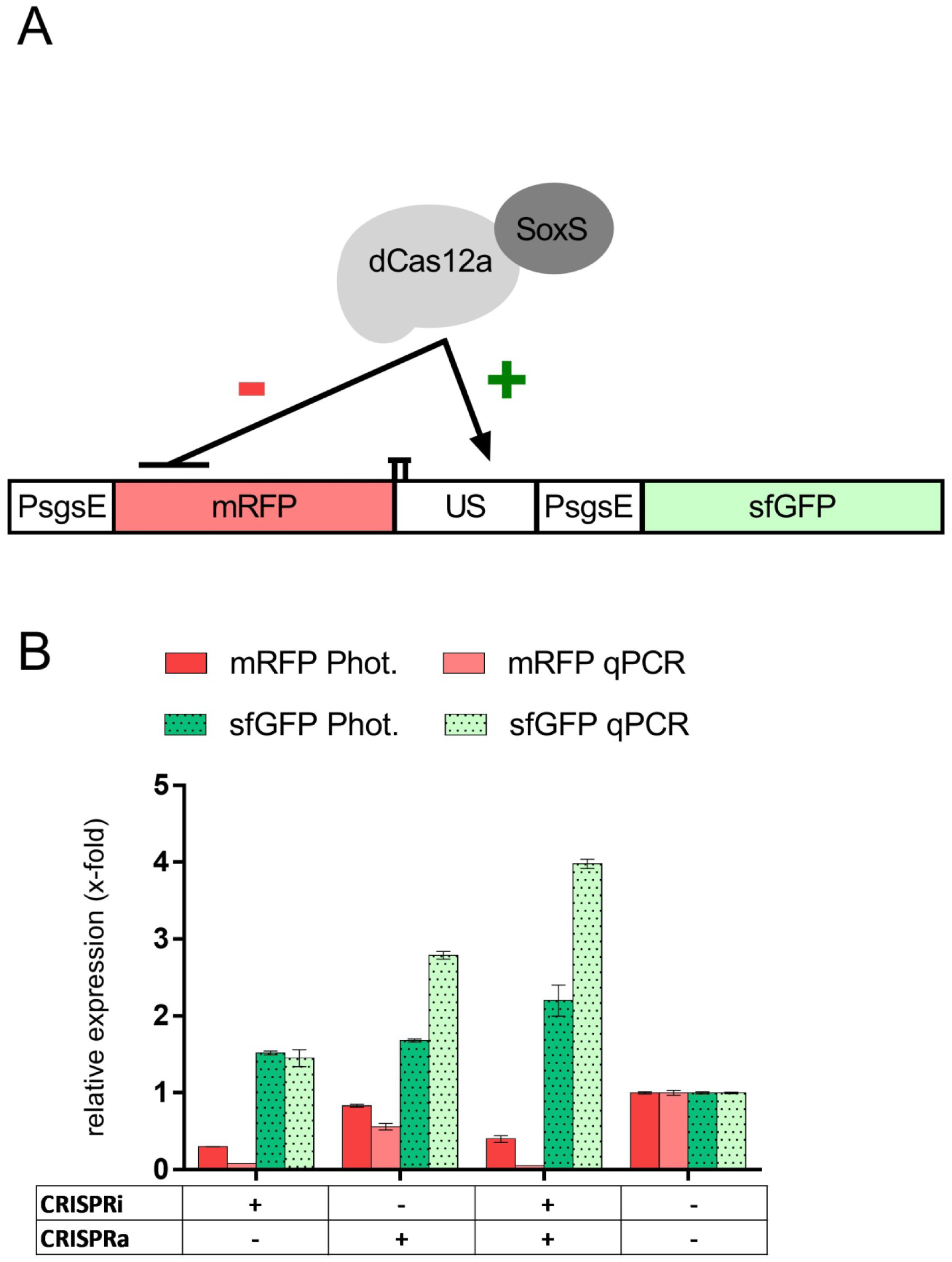
Simultaneous repression (CRISPRi, mRFP) and activation (CRISPRa, sfGFP) of fluorescence reporters. A) Schematic display of spacer sequences within the ORF of mRFP and in the upstream region (US) of PsgsE controlling the expression of sfGFP. B) Multiplex transcriptional perturbation was tested in *P. polymyxa* harboring a plasmid for the constitutive expression of GFP and mRFP fluorescence reporters. Single CRISPR arrays were designed targeting the ORF of mRFP or the upstream region of PsgsE controlling *sfgfp* expression (gRNA_a1). Expression of gRNAs resulted in the repression of mRFP or induction of sfGFP respectively. When both activated gRNAs were expressed simultaneously, obtained fluorescence results were similar to the expression of individual gRNAs alone. All results are depicted relative to an off-target CRISPR array encoding a spacer not present in the strain. Reporter expression was determined photometrically (after 48 cultivation) and by qPCR experiments (after 24 h cultivation) in biological triplicates.

### Multiplex CRISPRi to modify exopolysaccharide composition of *P. polymyxa* DSM 365

The most important advantage of Cas12a over the more commonly used Cas9 is the ability of the effector nuclease to process its own crRNA, allowing the simultaneous targeting of multiple loci through a single CRISPR-array^32^.

*P. polymyxa* DSM 365 is an avid producer of different exopolysaccharides. Depending on process conditions, variable polymer-mixtures are produced^39^. However, it has also been shown that the engineering of a polysaccharide structure with modified physicochemical properties is feasible^41^. The underlying gene cluster contains the two so-called initiating glycosyltransferases (GTi) PepC and PepQ, which are putatively responsible for the initiation of the biosynthesis of two distinct polysaccharides (Figure 4 A). Contrary to the first polymer (Paenan I), the second polymer (Paenan II) initiated by PepQ contains the deoxyhexose fucose (unpublished data). To evaluate the effects of CRISPRi constructs on genomic targets, the GTi PepQ was targeted alone or in combination with PepC. Strains harboring the plasmids pCRaiS (pCRai_soxS encoding an off-target spacer), pCRaiS_pepQ (targeting the ORF of *pepQ*) and pCRaiS_pepCQ (targeting the ORFs of *pepC* and *pepQ*) were constructed and used in EPS batch fermentations. Carbohydrate fingerprints of the obtained EPS were performed and compared to the respective knock-out strains (Figure 4 B).

**Figure 4:**
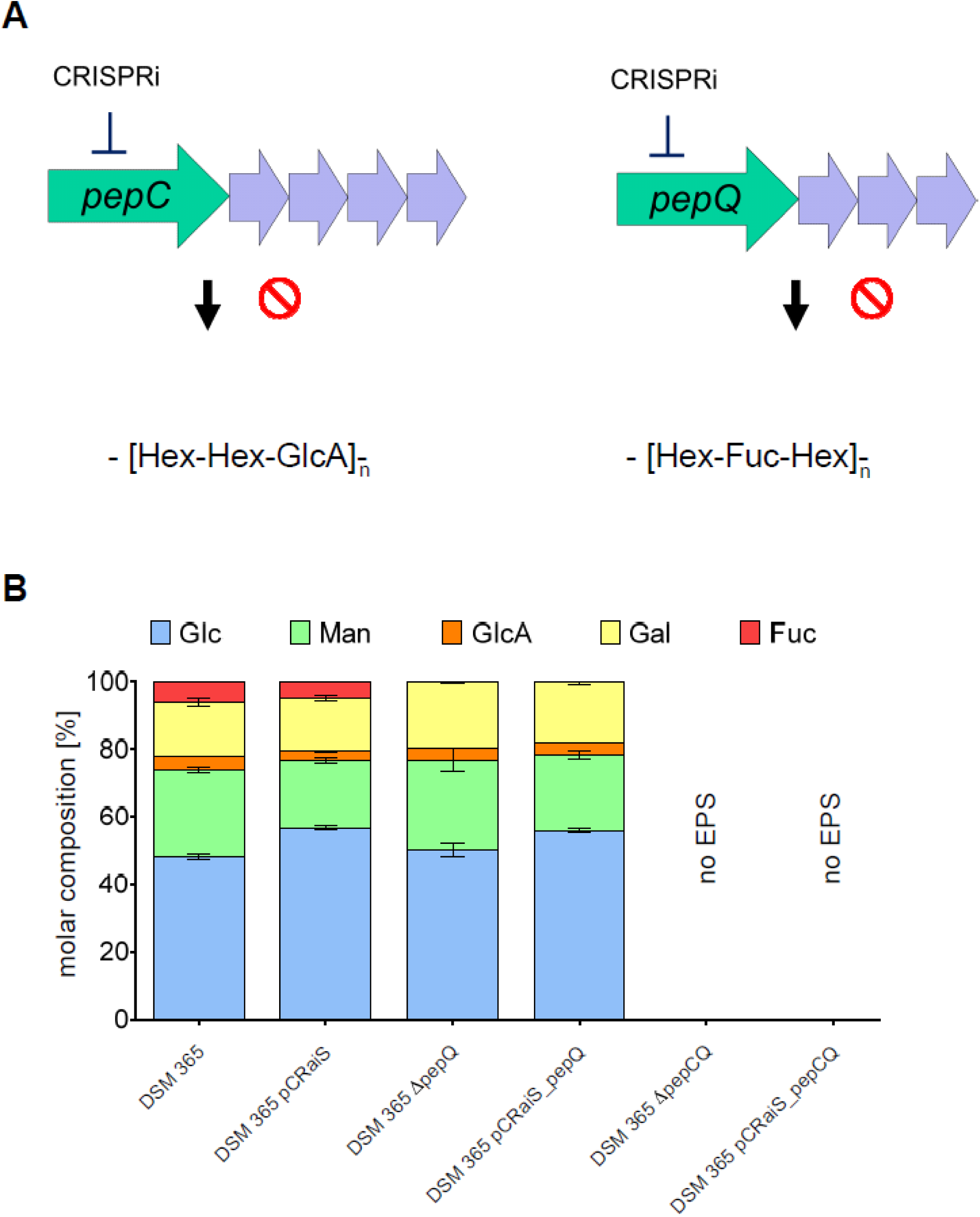
Carbohydrate fingerprint of the heteroexopolysaccharide of *P. polymyxa* DSM 365 and engineered variants. Transformation of *P. polymyxa* with a plasmid encoding the SoxS activator domain and off-target gRNA (pCRaiS) did not alter the EPS composition significantly. Expression of a gRNA targeting the ORF of the initiating glycosyltransferase *pepQ* (pCRaiS_pepQ) resulted in the loss of fucose within the EPS composition that was also observed in the KO strain *ΔpepQ*. Targeting the ORFs of both GTis (pCRaiS_pepCQ) did not yield any EPS resembling the same phenotype as the double KO *ΔpepCQ*, Δ: gene deletion by CRISPR-Cas9 mediated genome engineering. DSM 365: *P. polymyxa* DSM 365; pCRaiS: pCRai_soxS; Hex: hexose; GlcA: glucuronic acid; Fuc: fucose; Glc: glucose; Man: mannose; Gal: galactose

Strains harboring the pCRaiS plasmid expressing an off-target CRISPR-array did not show altered EPS composition. When the second GTi *pepQ* was targeted, fucose diminished, indicating the absence of Paenan II in the EPS mixture. In a next step, both initiating GTs were targeted simultaneously. With both GTis down-regulated, no EPS at all was produced. In order to evaluate whether the observed effects actually resulted from interference with the respective target genes, knock-out strains of the respective target genes with a previously established CRISPR-Cas9 genome editing system^41^ were constructed. Our experiments confirmed that the monomer composition was comparable to the respective CRISPRi variants (Figure 4 B). In the previous fluorescence reporter assays, reduced signal of mRFP and GFP respectively could still be detected in CRISPRi approaches. Contrary, for the glycosyltransferase targets, no significant differences in EPS composition were observed between the CRISPRi constructs and the knock-out strains, indicating a more efficient repression of the target genes in comparison to the fluorescence assays.

In conclusion, this approach showed that the developed pCRai_soxS tool can be used for the parallel screening for multiple interesting knock-out targets in the genome. In addition, the efficiency of this tool proved to be comparable to more laborious and time-consuming genome-editing approaches.

### Multiplex CRISPRi/a to screen for metabolic engineering targets of the butanediol biosynthesis pathway in *P. polymyxa* DSM 365

In a previous study we engineered the mixed acid pathway of *P. polymyxa* DSM 365 to increase the production of 2,3-BDL and remove undesirable side-products. Interestingly, knock-out of a lactate dehydrogenase (*ldh1*) resulted in an adapted growth behavior, increased biomass formation and consequently enhanced 2,3-BDL formation44. However, due to the presence of additional homologs of *ldh* within the genome, lactate formation could not be completely eliminated. Therefore, in order to demonstrate the capabilities of our dCas12a-based CRISPR-tool, all additional copies of *ldh* homologs were targeted in parallel. Leveraging the CRISPR-array processing abilities of Cas12a, each gene was targeted with two gRNAs at the same time. As decoupling of the 2,3-BDL biosynthesis from its natural regulon showed positive effects, the expression of the butanediol dehydrogenase should also be transcriptionally activated in *P. polymyxa* DSM 365 *Δldh1*. Therefore, the corresponding promoter was predicted using the Softberry CNNPromoter_b tool^54^. Three distinct gRNAs binding 106 - 180 bp upstream of the putative TSS were tested separately (bdh_a1 – a3). Thereby, three strains carrying plasmid constructs, each targeting 11 genomic sites in parallel were designed and evaluated in batch fermentations using microaerobic conditions (Figure 5 A). Due to the more stringent PAM requirements of Cas12a (TTTV), spacers more distant to the TSS had to be used for the *bdh* promoter compared to previous fluorescence experiments. 2,3-BDL fermentations were conducted for 72 h. Despite the fact that each ORF encoding the different lactate dehydrogenases was targeted with two gRNAs, lactate production in all strains was reduced only by ∼20 % compared to the strain harboring the off-target spacer (Figure 5 B). However, 2,3-BDL titers were increased from 27.5 g L^-1^ to 34.7 g L^-1^ for the *P. polymyxa* expressing the activated complex bdh_a3, corresponding to a 26 % increased product titer. Additionally, also the strains encoding bdh_a1 and bdh_a2 showed 25 % and 18 % increased 2,3-BDL titers respectively. All other end products of the mixed acid pathway that were not targeted by any gRNA remained similar in all variants (Figure 5, Figure S 4). Furthermore, 2,3-BDL yields were increased by approximately 20 % for all strains encoding target gRNAs, indicating a redirection of the carbon flux from lactate to 2,3-BDL (Table S4).

**Figure 5:**
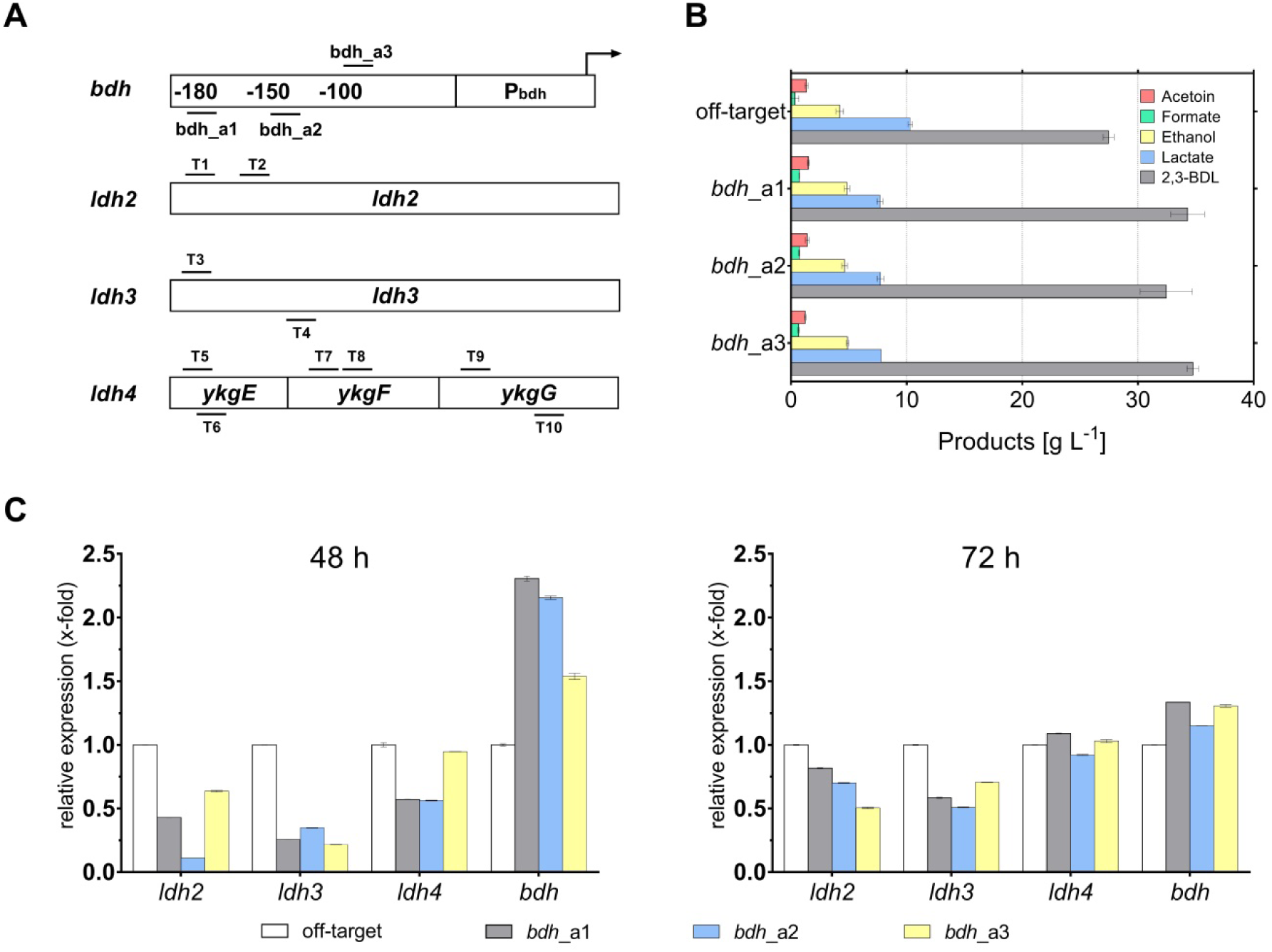
Multiplex CRISPRi and CRISPRa to engineer the mixed acid pathway of *P. polymyxa* and increase 2,3-BDL production. A) Schematic overview of gRNA binding sites. Three different gRNAs binding upstream of the P_bdh_ promoter were tested individually (bdh_a1-3). Simultaneously, all constructs also targeted three putative lactate dehydrogenase genes with a total of nine spacers (T1-T9) to knock-down the respective genes. B) Product titer obtained after 72 h cultivation at microaerobic conditions. Compared to an off-target construct, lactate production was reduced by ∼20 % in strains expressing target gRNAs. 2,3-BDL production was increased and reached a maximum of 34.7 g L^-1^ in the construct using bdh_a3 to target *bdh* expression. Depicted values represent the mean of biological duplicates. C) Expression of target genes was analyzed via qPCR after 48 h and 72 h of cultivation. After 48 h, transcription levels of lactate dehydrogenases (*ldh*2 - 4) were significantly reduced compared to a strain expressing off-target gRNAs. Furthermore, also expression of a butanediol dehydrogenase was increased. However, after 72 h of cultivation, effects of CRISPRi and CRISPRa were severely reduced.

While the general principle of our developed CRISPRi/a tool could be successfully demonstrated, effects of both transcriptional repression and activation were not as pronounced as observed in the fluorescence and EPS experiments. Expression analysis via qPCR revealed transcriptional perturbation of all *ldh* homologs ranging from 50 % to 80 % after 48 h of cultivation (Figure 5 C). Furthermore, two gRNAs binding 106 bp and 146 bp respectively upstream of the TSS of the *bdh* promoter caused more than a 2-fold increased expression on the transcriptional level. However, after 72 h effects on the transcriptional perturbation were significantly decreased. We hypothesize that the observed decreased effects by our dCas12a tool are a combined result of a rather low expression of the CRISPR-tool at microaerobic conditions and long cultivation times used for 2,3-BDL fermentations. Compared to other 2,3-BDL producing organisms such as *Serratia marcescens* or *Klebsiella pneumoniae, P. polymyxa* DSM 365 seems to be more sensitive towards high concentrations of 2,3-BDL with a toxic threshold at approximately 50 g L^-1 55^. As a result, using the applied microaerobic cultivation conditions, increased 2,3-BDL production is typically accompanied by elevated levels of lactate, which is used as a non-toxic, redox neutral electron acceptor^44^. Consequently, decreasing effects of transcriptional perturbation might be caused by genetic instabilities or by competing endogenous regulation mechanisms. Furthermore, the used spacers in a more distal region to the *bdh* promoter might be suboptimal for dCas12a-soxS. Due to the restrictive PAM site of Cas12a of *Acidaminococcus* sp., optimal distancing from the TSS might not always be possible and impede genome wide screenings. However, engineered variants of Cas12a have shown expanded binding motifs and enabled the targeting of otherwise inaccessible PAMs^36^.

The dynamic range of transcriptional perturbation described in this study was rather low. Depending on the applied conditions and targets, a 2 – 6.5 fold increased transcription was achieved by CRISPRa. While expression of some genes could be reduced by 80 % using CRISPRi, other targets were only poorly affected. Currently, each spacer requires to be tested separately to determine the efficiency of the system. Efficiencies of different PAMs have to be investigated more closely in the context of catalytically inactive Cas nucleases. Protein engineering approaches to modify dCas12a or the activator domain have previously demonstrated to be valid strategies to optimize and improve the binding efficiency of the effector module and dynamic output range for CRISPRa^23,36^.

## Conclusion

While CRISPRi has been continuously demonstrated in bacteria, CRISPRa technology is lacking behind on their eukaryotic counterparts. Currently available systems are still limited in their number of targets that can be modified in parallel due to the use of dCas9. In this study, we showed the first successful utilization of dCas12a for the simultaneous activation and repression of multiple genes in the alternative host organism *P. polymyxa* DSM 365.

While spacers particularly for CRISPRa still need to be optimized individually for each target promoter, utilization of SoxS as an activator domain enables more flexibility in the correct distancing to the target promoter compared to other tested activator domains. In this study, we demonstrated an efficient broad host range tool for the parallel transcriptional modulation of expression patterns in bacteria that can be applied for both metabolic engineering efforts and screening of potential targets for further studies. With ongoing studies using dCas12a-SoxS based tools for CRISPRa in bacterial hosts it will be possible to establish more precise design rule sets for the efficient positioning of the effector module for CRISPRi and CRISPRa to facilitate and accelerate the use of dCas12a based transcriptional perturbation tools.

Even though Cas12a is more restricted in terms of its PAM sequence compared to Cas9, its ability to efficiently process its own gRNAs makes it a promising tool to orchestrate sophisticated genetic reprogramming of bacterial cells or to screen for engineering targets in the genome.

In conclusion, this work demonstrated both, the simultaneous activation and repression of multiple targets in the genome of *P. polymyxa* using a single CRISPR array and represents therefore an important extension of current Cas9-based tools. We demonstrated that the developed tool is functional in common bacterial cell factories such as *E. coli* as well as in the Gram-positive alternative host organism *P. polymyxa*. Usage of multiplex transcriptional perturbation will facilitate the combinatorial regulation of complex pathways.

## Supporting information

Manuscript file

## Acknowledgement

This work was supported by the German Federal Ministry of Education and Research (BMBF) in frame of the project MaPolKo (number 03VP02560). MAGK would like to acknowledge funding from the U.S. National Science Foundation, award MCB-1817631.

## Supporting Information

Metabolic pathway scheme of butanediol production and polysaccharide synthesis, bacterial strains, plasmids, DNA and oligonucleotide sequences, plasmid map of pCRai, fluorescence reporter results for *E. coli* S17-1, fermentation profiles and product titers

