## Supplementary material for "A novel prokaryotic CRISPR-Cas12a based tool for programmable transcriptional activation and repression": Manuscript file

### Supporting Information

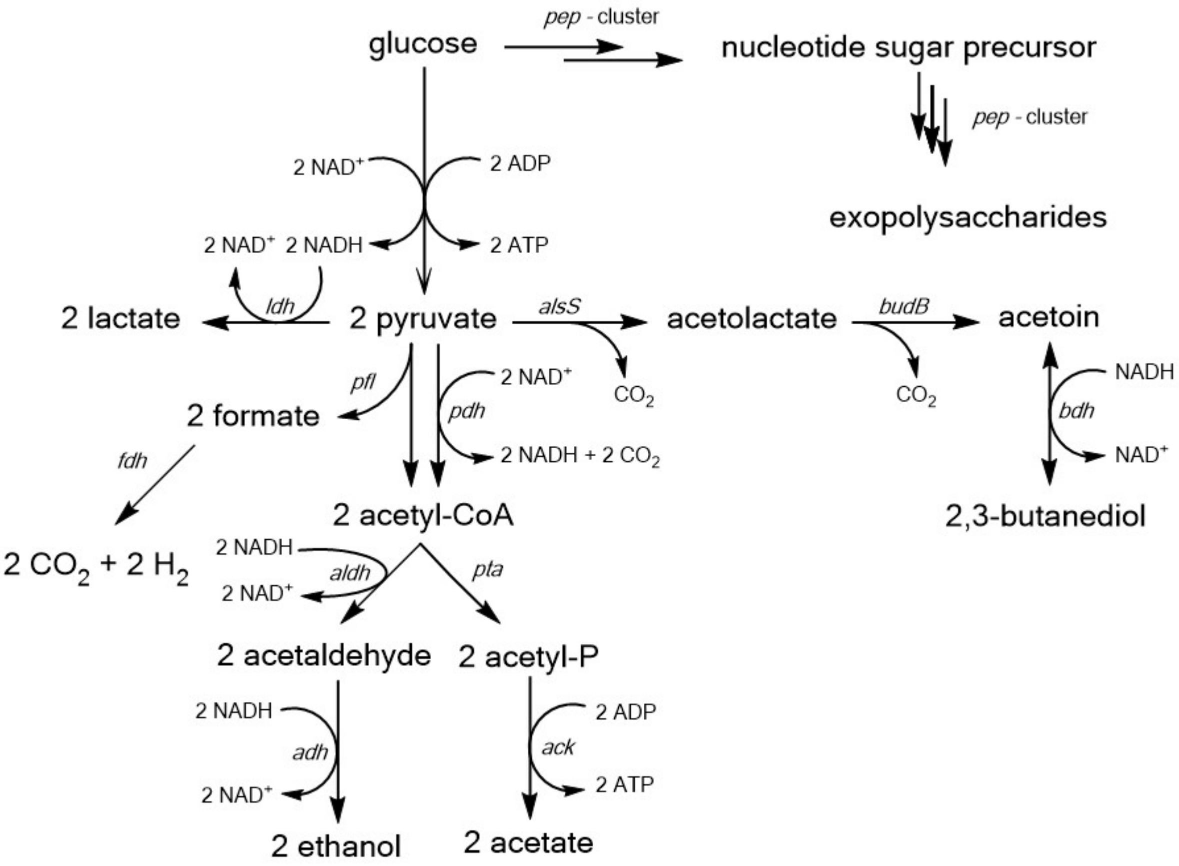


**Figure S 1: Overview of the mixed acid pathway and butanediol biosynthesis of *Paenibacillus polymyxa* in microaerobic conditions**. Two mols of NADH are formed during glycolysis that need to be regenerated in order to maintain redox balance. Only 1 mol NADH is converted to NAD+ in the 2,3-BDL pathway. Therefore, other redox neutral end products such as lactate or ethanol compete for the intermediate pyruvate. Furthermore, glucose is utilized for the production of exopolysaccharides (EPS). High molecular weight biopolymers are assembled by multiple glycosyltransferases encoded in the *pep*-cluster from nucleotide sugar precursor substrates. Adapted from Schilling et al., (2020)[1].

### Table S 1: Bacterial strains used in this study

| Bacterial Strains | Genotype | Reference |
|---|---|---|
| ***Escherichia coli* S17-1** | Conjugation strain; recA pro hsdR; RP42Tc::Mu-Km::Tn7 integrated into the chromosome | ATCC 47055 |
| ***P. polymyxa* DSM365** | wild type | DSMZ |
| ***P. polymyxa* DSM365 *Δldh1*** | DSM365 *Δldh1* | Schilling et al., (2020) |
| ***P. polymyxa* DSM365 *ΔpepCQ*** | DSM365 *ΔpepC* | This study |
| ***P. polymyxa* DSM365 *ΔpepQ*** | DSM365 *ΔpepQ* | This study |
| ***P. polymyxa* DSM365 *ΔpepCQ*** | DSM365 *ΔpepC ΔpepQ* | This study |

In addition to the strains listed above, each plasmid listed in Table S 2 was used for the transformation of *E. coli* S17-1 as well as for the conjugational transformation of *P. polymyxa* DSM 365.

### Table S 2: Plasmids used in this study

| Plasmids | Description | Reference |
|---|---|---|
| **pCasPP** | *P. polymyxa* CRISPR-Cas9 genome editing plasmid | Rütering et al (2017) |
| **pCasPPH_pepC** | pepC targeting knock out plasmid containing repair template | This study |
| **pCasPPH_pepQ** | pepQ targeting knock out plasmid containing repair template | This study |
| **pCRai** | PsgsE-AsdCas12a; Pgapdh-off target gRNA; neo; oriT, repU | This study |
| **pCRaiGFP** | PsgsE-AsdCas12a; Pgapdh-off target gRNA;neo; oriT, repU; PsgsE-sfGFP | This study |
| **pCRaiGmR_soxS** | PsgsE-AsdCas12a-soxS; Pgapdh-off target gRNA; neo; oriT, repU; PsgsE_mRFP-US-PsgsE-sfGFP | This study |
| **pCRaiGFP_GltC** | Test plasmid for CRISPRa targeting US region of PsgsE-sfGFP - off target gRNA | This study |
| **pCRaiGFP_GltC_a1** | Test plasmid for CRISPRa targeting US region of PsgsE-sfGFP - sgRNA_a1 | This study |
| **pCRaiGFP_GltC_a2** | Test plasmid for CRISPRa targeting US region of PsgsE-sfGFP - sgRNA_a2 | This study |
| **pCRaiGFP_GltC_a3** | Test plasmid for CRISPRa targeting US region of PsgsE-sfGFP - sgRNA_a3 | This study |
| **pCRaiGFP_GltC_a4** | Test plasmid for CRISPRa targeting US region of PsgsE-sfGFP - sgRNA_a4 | This study |
| **pCRaiGFP_sig70** | Test plasmid for CRISPRa targeting US region of PsgsE-sfGFP - off target gRNA | This study |
| **pCRaiGFP_sig70_a1** | Test plasmid for CRISPRa targeting US region of PsgsE-sfGFP - sgRNA_a1 | This study |
| **pCRaiGFP_sig70_a2** | Test plasmid for CRISPRa targeting US region of PsgsE-sfGFP - sgRNA_a2 | This study |
| **pCRaiGFP_sig70_a3** | Test plasmid for CRISPRa targeting US region of PsgsE-sfGFP - sgRNA_a3 | This study |

| | | |
|---|---|---|
| **pCRaiGFP_sig70_a4** | Test plasmid for CRISPRa targeting US region of PsgsE-sfGFP - sgRNA_a4 | This study |
| **pCRai_GFP_rpoD_a1** | Test plasmid for CRISPRa targeting US region of PsgsE-sfGFP - sgRNA_a1 | This study |
| **pCRai_GFP_rpoD_a2** | Test plasmid for CRISPRa targeting US region of PsgsE-sfGFP - sgRNA_a2 | This study |
| **pCRai_GFP_rpoD_a3** | Test plasmid for CRISPRa targeting US region of PsgsE-sfGFP - sgRNA_a3 | This study |
| **pCRai_GFP_rpoD_a4** | Test plasmid for CRISPRa targeting US region of PsgsE-sfGFP - sgRNA_a4 | This study |
| **pCRai_GFP_rpoD** | Test plasmid for CRISPRa targeting US region of PsgsE-sfGFP - off target gRNA | This study |
| **pCRai_GFP_soxS_a1** | Test plasmid for CRISPRa targeting US region of PsgsE-sfGFP - sgRNA_a1 | This study |
| **pCRai_GFP_soxS_a2** | Test plasmid for CRISPRa targeting US region of PsgsE-sfGFP - sgRNA_a2 | This study |
| **pCRai_GFP_soxS_a3** | Test plasmid for CRISPRa targeting US region of PsgsE-sfGFP - sgRNA_a3 | This study |
| **pCRai_GFP_soxS_a4** | Test plasmid for CRISPRa targeting US region of PsgsE-sfGFP - sgRNA_a4 | This study |
| **pCRai_GFP_soxS** | Test plasmid for CRISPRa targeting US region of PsgsE-sfGFP - off target gRNA | This study |
| **pCRai_GFP_CRP_a1** | Test plasmid for CRISPRa targeting US region of PsgsE-sfGFP - sgRNA_a1 | This study |
| **pCRai_GFP_CRP_a2** | Test plasmid for CRISPRa targeting US region of PsgsE-sfGFP - sgRNA_a2 | This study |
| **pCRai_GFP_CRP_a3** | Test plasmid for CRISPRa targeting US region of PsgsE-sfGFP - sgRNA_a3 | This study |
| **pCRai_GFP_CRP_a4** | Test plasmid for CRISPRa targeting US region of PsgsE-sfGFP - sgRNA_a4 | This study |
| **pCRai_GFP_CRP** | Test plasmid for CRISPRa targeting US region of PsgsE-sfGFP - off target gRNA | This study |
| **pCRaiGFP_soxS_sfGFP_T1** | Test plasmid for sfGFP repression sgRNA_T1 | This study |
| **pCRaiGFP_soxS_sfGFP_T2** | Test plasmid for sfGFP repression sgRNA_T2 | This study |
| **pCRaiGFP_soxS_sfGFP_T3** | Test plasmid for sfGFP repression sgRNA_T3 | This study |
| **pCRai_pepQ_T1** | Plasmid for repression of pepQ sgRNA T1 | This study |
| **pCRai_pepCQ_T1** | Plasmid for dual repression of pepQ and pepC | This study |
| **pCRai_soxS_ldhmultibdh1** | Multiplex knock-down of repression of ldh2, ldh3, ldh4 and activation of Pbdh (sgRNA 1) | This study |
| **pCRai_soxS_ldhmultibdh2** | Multiplex knock-down of repression of ldh2, ldh3, ldh4 and activation of Pbdh (sgRNA 2) | This study |
| **pCRai_soxS_ldhmultibdh3** | Multiplex knock-down of repression of ldh2, ldh3, ldh4 and activation of Pbdh (sgRNA 3) | This study |
| **pCRaiGmR_soxS_mT2** | PsgsE-mRFP-US-PsgsE_sfGFP expression plasmid with gRNA for mRFP repression (sgRNA T2) | This study |
| **pCRaiGmR_soxS_dualT2** | PsgsE-mRFP-US-PsgsE_sfGFP expression plasmid with gRNA for mRFP repression (sgRNA T2) and GFP activation (sgRNA a1) | This study |
| **pCRaiGmR_soxS_GFPa1** | PsgsE-mRFP-US-PsgsE_sfGFP expression plasmid with gRNA and GFP activation (sgRNA a1) | This study |

**Table S 3: Oligonucleotides and primers used in this study.** Overhangs used for Golden Gate Assembly or Gibson isothermal assembly are depicted in lower case. Restriction sites are underlined.

| Name | SEQUENCE 5'->3' | Comment |
|---|---|---|
| **soxS_fw** | ATGTTGGTCTCAACGCTGATGTCCCATCAGAAAATTATTCAGGATC | Cloning of activator domain |
| **soxS_rev** | TAAAAGGTCTCGTACCATTTACAGGCGGTGGCGATAATCG | Cloning of activator domain |
| **sig70_fw** | ATGTTGGTCTCAACGCTGATGATAGAGGACACACCATTCCGT | Cloning of activator domain |
| **sig70_rev** | TAAAAGGTCTCGTACCATTCATTGCCATTCCTCCTCCTTCC | Cloning of activator domain |
| **cAMP_fw** | ATGTTGGTCTCAACGCTGATGCATCCCTATTATCGTTCATTAGAAAAAA | Cloning of activator domain |
| **cAMP_rev** | TAAAAGGTCTCGTACCATTTAATGCGGATCACTACGCAGCAA | Cloning of activator domain |
| **GltC_fw** | ATGTTGGTCTCAACGCTGGTGGAATTACGACAGTTACAATACTTCATGAA | Cloning of activator domain |
| **GltC_rev** | TAAAAGGTCTCGTACCATTCACTGCCCTAAATCCGCTTTTG | Cloning of activator domain |
| **RpoD_fw** | ATGTTGGTCTCAACGCTGGTGCTGTATCCTTCCATTGATGAAATG | Cloning of activator domain |
| **RpoD_rev** | TAAAAGGTCTCGTACCATTTATTCTTCTTCGCCGCTTTCAAGT | Cloning of activator domain |
| **As_CRISPR_fw** | GGGGATACGCTAATTTCTACTCTTGTAGATAAGTCTTCTCAGCCG | Construction of pCRai |
| **As_CRISPR_rev** | AAATCCAGATGGAGTATGTCTTCACCGGTGGAAAGCG | Construction of pCRai |
| **AsCRISPR_vec_fw** | CACCGGTGAAGACATACTCCATCTGGATTTGTTCAGAACGC | Construction of pCRai |
| **AsCRISPR_vec_rev** | GTAGAAATTAGCGTATCCCCTTTCAGATACTCGCAC | Construction of pCRai |
| **enAsCPF1_fw** | TTAGGCTTTTACTTAATGACACAGTTTGAAGGCTTTACGAATCTG | Construction of pCRai |
| **enAsCPF1_rev** | CGTAGATCTGAATTCTTATTACCCGAGACCTACCCAATGCG | Construction of pCRai |
| **enAsCPF1_Vector.fw** | GGTCTCGGGTAATAAGAATTCAGATCTACGCGTTCCCCGC | Construction of pCRai |
| **enAsCPF1_Vector.REV** | TTCAAACTGTGTCATTAAGTAAAAGCCTAAAATCCCCCTTCGTT | Construction of pCRai |
| **GFP_vecint_rev** | gcaacgcggcctttttacggTTCCTGGCCATATGACGATCCTCCTTACCTCTCATTG | sfGFP plasmid cloning |
| **GFP_vecint_fw** | gcccgcaaccggcgcatcaagcccgccGACTAGCAATATGAAACACGGAAAAAATCAAGC | sfGFP plasmid cloning |
| **sgsE-a1_fw** | agatTTTATCTCACATAATAGGGCT | Test gRNA for activation of PsgsE-sfGFP |
| **sgsE-a1_rev** | gagtAGCCCTATTATGTGagatAA | Test gRNA for activation of PsgsE-sfGFP |
| **sgsE-a2_fw** | agatAGTCTATATCAATCGGTAAC | Test gRNA for activation of PsgsE-sfGFP |

| | | |
|---|---|---|
| **sgsE-a2_rev** | gagtGTTACCGATTGATATAGACT | Test gRNA for activation of PsgsE-sfGFP |
| **sgsE-a3_fw** | agatATCCTCATATTTTCCTAGTA | Test gRNA for activation of PsgsE-sfGFP |
| **sgsE-a3_rev** | gagtTACTAGGAAAATATGAGGAT | Test gRNA for activation of PsgsE-sfGFP |
| **sgsE-a4_fw** | agatCATCGATGGCGACATTGATA | Test gRNA for activation of PsgsE-sfGFP |
| **sgsE-a4_rev** | gagtTATCAATGTCGCCATCGATG | Test gRNA for activation of PsgsE-sfGFP |
| **sfGFP1_T1_fw** | agatCGTGCGTGGCGAGGGTGAAG | Test gRNA for repression of PsgsE-sfGFP |
| **sfGFP1_T1_rev** | gagtCTTCACCCTCGCCACGCACG | Test gRNA for repression of PsgsE-sfGFP |
| **sfGFP_T2_fw** | agatCCATTAGTTGCGTCACCTTC | Test gRNA for repression of PsgsE-sfGFP |
| **sfGFP_T2_rev** | gagtGAAGGTGACGCAACTAATGG | Test gRNA for repression of PsgsE-sfGFP |
| **sfGFP_T3_fw** | agatAGCTCAATGCGGTTTACCAG | Test gRNA for repression of PsgsE-sfGFP |
| **sfGFP_T3_rev** | gagtCTGGTAAACCGCATTGAGCT | Test gRNA for repression of PsgsE-sfGFP |
| **mRFP_T_fw** | agatAAAGTTCGTATGGAAGGTTC | gRNA for repression of PsgsE-mRFP |
| **mRFP_T_rev** | gagtGAACCTTCCATACGAACTTT | gRNA for repression of PsgsE-mRFP |
| **sfGFP_dual_fw** | agatTTATCTCACATAATAGGGCTTAATTTCTACTC | Test gRNA for repression of PsgsE-mRFP and simultaneous activation of PsgsE-sfGFP |
| **sfGFP_dual_rev** | acaaGAGTAGAAATTAAGCCCTATTATGTGAGATAA | Test gRNA for repression of PsgsE-mRFP and simultaneous activation of PsgsE-sfGFP |
| **mRFP_dual_fw** | ttgtAGATAAAGTTCGTATGGAAGGTTC | Test gRNA for repression of PsgsE-mRFP and simultaneous activation of PsgsE-sfGFP |
| **mRFP_dual_rev** | gagtGAACCTTCCATACGAACTTTATCT | Test gRNA for repression of PsgsE-mRFP and simultaneous activation of PsgsE-sfGFP |
| **GGA_sfGFP_fw** | AATT<u>GGTCTC</u>AAACTCAATATGAAACACGGAAAAAATCAAGCAG | Cloning of PsgsE-mRFP-US-PsgsE-sfGFP expression cassette |
| **GGA_sfGFP_rev** | AATT<u>GGTCTC</u>ACTAGTATGACGATCCTCCTTACCTCTCATTG | Cloning of PsgsE-mRFP-US-PsgsE-sfGFP expression cassette |
| **GGA_mRFP_fw** | AATT<u>GGTCTC</u>ATCGACCTGCATACTAGCCTGTTACAGGCATATTCATATCAATGTC | Cloning of PsgsE-mRFP-US-PsgsE-sfGFP expression cassette |
| **GGA_mRFP_rev** | CTTT<u>GGTCTC</u>AAGTTAAGCACCGGTGgagtGACGAC | Cloning of PsgsE-mRFP- |

| | | |
|---|---|---|
| | | US-PsgsE-sfGFP expression cassette |
| **pepQ_T1 fw** | agatCACCAGCAGTGCAGACAATC | gRNA for repression of pepQ |
| **pepQ_T1_rev** | gagtGATTGTCTGCACTGCTGGTG | gRNA for repression of pepQ |
| **pepCQ_T1_fw** | agatCCTGACATGACAGCCTCGCCTAATTTCTACT | gRNAs for dual repression of pepCQ |
| **pepCQ_T1_rev** | caagAGTAGAAATTAGGCGAGGCTGTCATGTCAGG | gRNAs for dual repression of pepCQ |
| **pepCQ2_T1_fw** | cttgTAGATCACCAGCAGTGCAGACAATC | gRNAs for dual repression of pepCQ |
| **pepCQ_T1_rev** | gagtGATTGTCTGCACTGCTGGTGATCTA | gRNAs for dual repression of pepCQ |
| **bdh1_fw** | cttgTAGATGACACTCATTCTGTGGTATA | gRNA cloning for Pbdh activation |
| **bdh1_rev** | gagtTATACCACAGAATGAGTGTCATCTA | gRNA cloning for Pbdh activation |
| **bdh2_fw** | cttgTAGATTTCTTGTCTTTGCTTCAATT | gRNA cloning for Pbdh activation |
| **bdh2_rev** | gagtAATTGAAGCAAAGACAAGAAATCTA | gRNA cloning for Pbdh activation |
| **bdh3_fw** | gagtGCTCGTTACTTTTATACAAA | gRNA cloning for Pbdh activation |
| **bdh3_rev** | gagtTTTGTATAAAAGTAACGAGCATCTA | gRNA cloning for Pbdh activation |
| **sfGFP_qPCR_fw** | CCCTATTCTGGTGGAACTGGATGG | qPCR primer |
| **sfGFP_qPCR_rev** | CAGTAGTACAGATGAACTTCAGCGTC | qPCR primer |
| **gyrA_qPCR_fw** | GAGATATGGCCGCTGCGATG | qPCR primer |
| **gyrA_qPCR_rev** | GCTCTCTTCACCATCGTAGTTCGG | qPCR primer |
| **mRFP_qPCR_fw** | TACCTGAAACTGTCCTTCCCGG | qPCR primer |
| **mRFP_qPCR_rev** | GTAGATGAACTCACCGTCTTGCAGG | qPCR primer |
| **ldh3_qPCR_fw** | GAGTCATTGGATCA GA CGTTGC | qPCR primer |
| **ldh3_qPCR_rev** | AACTCGGAGTCACCGTGTTCTCC | qPCR primer |
| **ldh2_qPCR_fw** | GATTATCGGGGTTGGACGCATTGG | qPCR primer |
| **ldh2_qPCR_rev** | CAAGGTTGTGAATGGAATGACTTCCTTG | qPCR primer |
| **ldh4_qPCR_fw** | GAATGAGCCGGACCTGGATGCG | qPCR primer |
| **ldh4_qPCR_rev** | CAATCCTCGACTGGATCGGTGTTC | qPCR primer |
| **bdh_qPCR_fw** | AGATGCTCAGCATCCATTGACTGG | qPCR primer |
| **bdh_qPCR_rev** | CAACGACACGGTCGCCAACCTG | qPCR primer |

**Table S 4: spacer sequences and protospacer adjacent motifs (PAM) of each gRNA used in this study**

| Target | Spacer 5’-3’ | PAM 5’-3’ |
|---|---|---|
| sgsE-a1 | TTTATCTCACATAATAGGGCT | TTTC |
| sgsE-a2 | AGTCTATATCAATCGGTAAC | TTTC |
| sgsE-a3 | ATCCTCATATTTTCCTAGTA | TTTC |
| sgsE-a4 | CATCGATGGCGACATTGATA | TTTC |
| sfGFP_T1 | CGTGCGTGGCGAGGGTGAAG | TTTC |
| sfGFP_T2 | CCATTAGTTGCGTCACCTTC | TTTA |
| sfGFP_T3 | AGCTCAATGCGGTTTACCAG | TTTC |
| mRFP | AAAGTTCGTATGGAAGGTTC | TTTC |
| pepQ_ | CACCAGCAGTGCAGACAATC | TTTC |
| pepC | CCTGACATGACAGCCTCGCC | TTTA |
| bdh1 | TGACACTCATTCTGTGGTAT | TTTA |
| bdh2 | TTTCTTGTCTTTGCTTCAATT | TTTC |
| bdh3 | GCTCGTTACTTTTATACAAA | TTTG |
| ldh2_T1 | CGATTGGCAGCACAGTTTCC | TTTC |
| ldh2_T2 | CAGTTATGGATCTCTTTGCT | TTTA |
| ldh3_T1 | ATCACAAATGGACTGATTAA | TTTC |
| ldh3_T2 | TACCATATACGTTACAATGT | TTTG |
| ykgE_T1 | ATCCCAATAGCCGCTGTTAT | TTTC |
| ykgE_T2 | TTCCAGCCTCCGTGCTGAAT | TTTG |
| YkgF_T1 | CGTTACGCAAACGCTCTGTA | TTTC |
| ykgF_T2 | TACAAATAGATTTAAGTAAT | TTTC |
| ykgG_T1 | ATGAAGCCACGCTTCATGGG | TTTG |
| ykgG_T2 | TGCGCGATCCAGTCTGCGGT | TTTG |
| off-target | AAGTCTTCTCAGCCGCTACA | TTTV |

**Table S 5: nucleotide sequences of activator domains used in this study**

| Activator | Protein Class | Nucleotide sequence (5’-3’) |
|---|---|---|
| SoxS<br>(324 bp) | AraC/XylS family | ATGTCCCATCAGAAAATTATTCAGGATCTTATCGCATGGATTGACGAGCAT<br>ATTGACCAGCCGCTTAACATTGATGTAGTCGCAAAAAAATCAGGCTATTCA<br>AAGTGGTACTTGCAACGAATGTTCCGCACGGTGACGCATCAGACGCTTGG<br>CGATTACATTCGCCAACGCCGCCTGTTACTGGCCGCCGTTGAGTTGCGCAC<br>CACCGAGCGTCCGATTTTTGATATCGCAATGGACCTGGGTTATGTCTCGCA<br>GCAGACCTTCTCCCGCGTTTTCCGTCGGCAGTTTGATCGCACTCCCAGCGA<br>TTATCGCCACCGCCTGTAA |
| GltC<br>(939 bp) | LysR family | GTGGAATTACGACAGTTACAATACTTCATGAAGGTAGCGCAAAAAGAACA<br>TGTCACTCAGGCAGCCGAGGAGCTACATGTGGCGCAATCTGCGGTAAGTC<br>GGCAAATTCATCAGCTTGAAGAGGAGCTGGGAGTTAATCTTTTTATGCAA<br>AAGGGTCGCAATTTGCAGCTTACGCCAGTGGGACAGCTATTTTGCAAACG<br>GGTTGAGACGATAATAAAAGATCTAGAGCGAGCTGTTTTGGAGGTTCATG<br>AGTTTTTGGATCCTGAGGGGGGCGAAATCCGCATCGGTTTTCCGCACAGT<br>TTGGGGATTCATCTTATTCCTACGGTAGTGGCTGAATTCCGAAAGCGGTAT<br>CCCAATGTTAAATTTAGATTTAAACAGGGGATGTATCCGAGTTTAATTCGT<br>GATGTGTTAGCGGGTGAGGTGGACTTGGCTTTTGTATCTCCTTTTCCCGAT<br>CGTCATGATCATGTGGAGGGGGATGTGGTGCTGACCGAGGAGTTGTTTGC<br>TGTTCTCCCGCCCAATCATCCTTTAGCTACGGCCAAACATATCACGCTGAGT<br>CAGCTTAAAGGAGAAAAATTTATTCTGTTCAGAGATGGATATTCGTTACGT |

| | | |
|---|---|---|
| | | CCGATTGTCTGGCAGGCTTGTCTGGAGGCAGGATTTACGCCAGATATCGC<br>TTTTGAAGGTGAAGAGACAGATACGATTCGCGGATTAGTTGCGGCTGGTA<br>TGGGCGTTAGCCTTTTGCCTGAAATGGCGTTGTTTCAGACCAACCCGCTTC<br>AACCTGCAAGAGTGTCAATTGTTGATCCAGAGGTCACGCGAACTATAGGC<br>TTGATTCATCGCAAAGATGACAAGCTACCATTAGTTGCAAAGTCATTCCGG<br>ACTTTTTTGTTGCAATATTTTGGTCTTAAAGGTAGTGTTGCTACTCCGAATG<br>GAAGTGCAAAAGCGGATTTAGGGCAGTGA |
| RpoD<br>(666 bp) | Sigma factor | ATGATAGAGGACACACCATTCCGTCAGGAGGGTGTGTCCTTTTTAAATTAT<br>TTTTTCGAAATATGCAGGGGAAGGAAATTTCCTGGCGTCTATAGTATTGG<br>GAAGGGGGAGAGCGTCATTGATATAGAGAGAATCATTGAACGAATACGG<br>TCTAGTGACCGGCAGGCGTTCCGTGAGATCGTAGAGCTGTACAGCAAGCA<br>TGTATTTCATATCGCCTATTCGGTGCTGCATGACAGCAAGGAAGCAGAAG<br>ATGCCGCGCAGGAGGCATTCGTTCAGGTGTACAAGTCTCTCCCCCAGTACC<br>GGAACGAAGGTTTTAAAACGTGGCTGAGCCGAATTGCGCTCCACAAAGCG<br>CTGGATATTAAGCGTAAACAGGACAGACGGCCCGCAGAGCTAATAGACGT<br>CGCACAATCCCTTGTACAGCTCCCTTCCCGGGATGAAGATGTGCTGGCTCG<br>TCTCATCCGCGAGGAGCAAATTAAAGAACTATCACAAAAAATCGCTCAGTT<br>GCCCCAGCAACATCGGGATATTATTCAAGCCTACTATATGCAGGGCAAAA<br>CCTACGACCAAATTGCAGAAGAGACGCAGGTAGCTTTAAAAACGGTAGAG<br>TCGAGACTCTACCGGGCGCGACTATGGATTCGAAACCACTGGAAGGAGG<br>AGGAATGGCAATGA |
| RpoZ<br>(204 bp) | DNA directed<br>RNA<br>polymerase<br>subunit | GTGCTGTATCCTTCCATTGATGAAATGATGAAAAAGGCAGATAGTAAGTA<br>TTCGTTGGTTGTAGCGGCTTCACGTCGTGCCAGACAGTTGCGCGAGGGTG<br>AAAAGACGACCCTCAAGAATCCTAAATCCCACAAACAGGTTGGAGTAGCT<br>TTGGAAGAGATCTATGCAGATCATCTTCTACTTGAAAGCGGCGAAGAAGA<br>ATAA |
| CRP<br>(582 bp) | cAMP<br>receptor | ATGCATCCCTATTATCGTTCATTAGAAAAAATGGCTCATCATGCCTCGATTA<br>TAGAGCAGGATTGGAACAGGTTTGTAGAGCAAACCCGTGTGAAGCATGTT<br>CAGCAGGGAACGGTTCTCATTGAAACGGGTGAGCCTGTGAAACATGCCTA<br>TTTTTGTTCAGAGGGATTATTCCGTTTATTTTATACATTAGTAGATGGCAGA<br>GAATATAATGTTGGTTTTTCGCCAGAAGATGATTACGTAACCTCTTATGGG<br>GCTATGATCCAAGGAGAACCATCCACTTTTTCGATTGAGGCCATGGAAGAT<br>TCGGTTGTCATTGAAATTCCATATCACGTATTGAAGGAGCTTATGGATACA<br>AGCCATATGTGGGAAAGATTCGTCCGTAAAAGTGTAGAAAGACTTTATAT<br>TCGAAAAGAGGAACGGGAGCGCGAATTATTGTATCTTTCCGCTAAGGAGC<br>GTTATCATGCTTTTCTTCTCAAATACCCCGGATTGGACAAACGAGTTGCCCA<br>ATATCATATAGCTTCGTACATAGGCGTCTCCCCTGTATCACTTAGTCGTTTG<br>CTGCGTAGTGATCCGCATTAA |

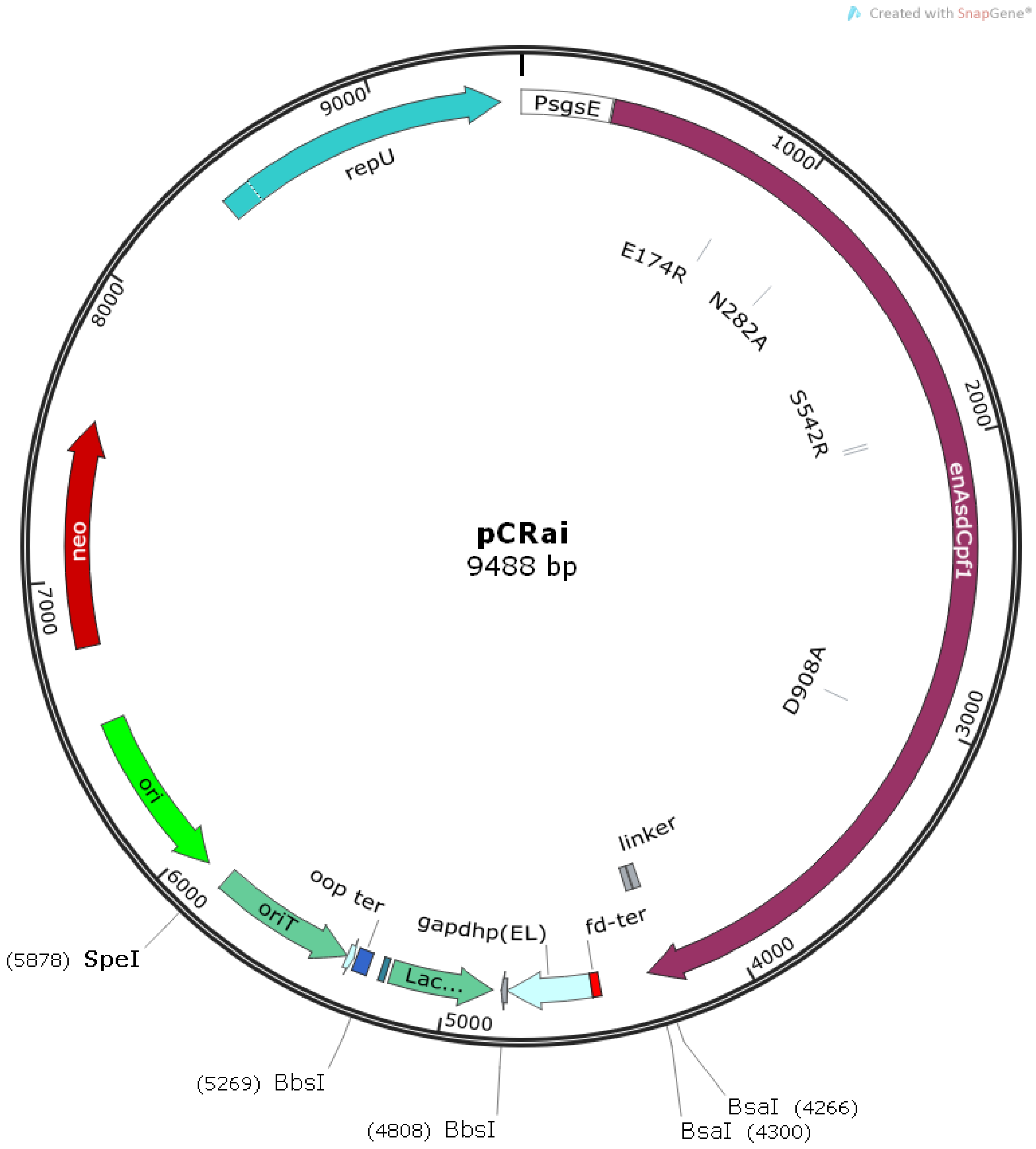


**Figure S 2: Plasmid map of pCRai.** The conjugational plasmid based on the pCasPP[2] construct was cloned by Gibson isothermal Assembly by replacing the Cas9-gRNA expression cassette with an engineered catalytically inactive variant of Cas12a[3] (E174R, N282A, S542R, K548R, D908A) and an appropriate CRISPR-array. Variable activator domains can be cloned by replacing a BsaI-cassette linked via a 10 aa flexible linker. gRNAs can be cloned by Golden Gate Assembly using BbsI by replacing a *lacZ* expression cassette.

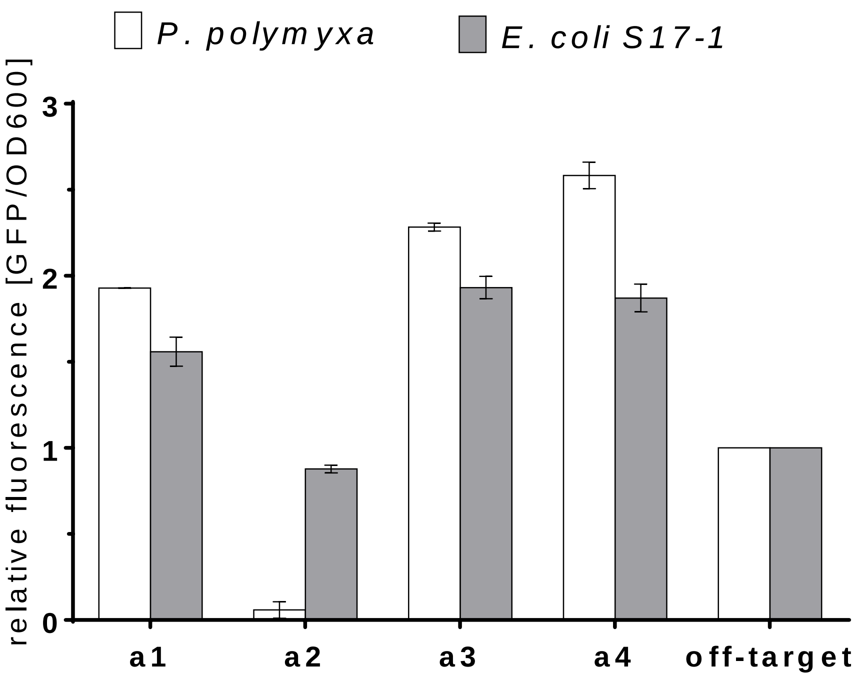


**Figure S 3: Relative fluorescence levels of *P. polymyxa* and *E. coli* S17-1 harboring pCRaiGFP_soxS variants.** 4 different gRNAs (a1-a4) were expressed and normalized fluorescence levels (Ex. 488 nm Em. 515 nm) were compared to a strain expressing an off-target spacer. Spacers a1, a3 and a4 showed increased GFP fluorescence in both *P. polymyxa* and *E. coli* S17-1 demonstrating similar effects in both, Gram-positive and Gram-negative host organisms.

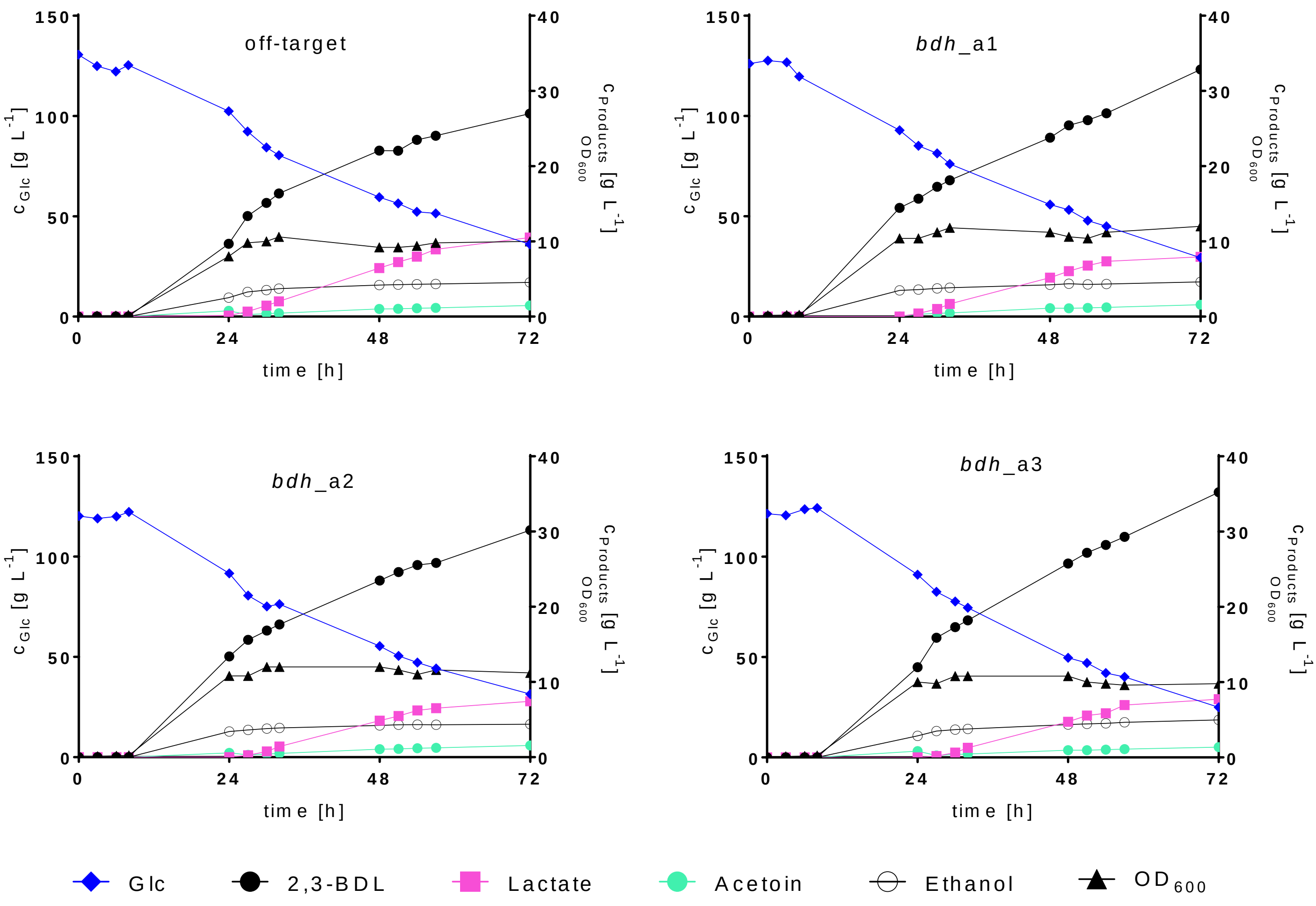


**Figure S 4: Fermentation profile of 2,3-BDL fermentations.** Overview of fermentation profiles of single batch fermentations. *P. polymyxa* DSM 365 was transformed with pCRai_soxS encoding either off-target gRNAs or a multiplex CRISPR-array targeting the open reading frame of three lactate dehydrogenases and the upstream region of $P_{bdh}$ with three gRNAs (bdh_a1-a3) individually. Lactate production was reduced by ~ 20 % in all variants expressing the corresponding gRNAs. Glc: Glucose; 2,3-BDL: 2,3-*R,R*-butantediol

**Table S 5 Final product titers and 2,3-BDL yield ($g_{BDL}$ $g_{Glc}^{-1}$) of 2,3-BDL fermentations.** *P. polymyxa* DSM 365 was transformed with pCRai_soxS encoding either off-target gRNAs or a multiplex CRISPR-array targeting the open reading frame of three lactate dehydrogenases and the upstream region of $P_{bdh}$ with three gRNAs (bdh_a1-a3) individually**.** Values represent mean and deviation of biological duplicates after 72 h cultivation in microaerobic conditions.

| | **2,3-BDL** | **Ethanol** | **Acetoin** | **Lactate** | **Formate** | **$Y_{PS}$** |
|---|---|---|---|---|---|---|
| **off-target** | 27.46 ± 0.47 | 4.21 ± 0.33 | 1.34 ± 0.14 | 10.30 ± 0.20 | 0.34 ± 0.34 | 0.29 ± 0.01 |
| **bdh_a1** | 34.30 ± 1.45 | 4.84 ± 0.23 | 1.50 ± 0.08 | 7.69 ± 0.26 | 0.72 ± 0.03 | 0.34 ± 0.00 |
| **bdh_a2** | 32.44 ± 2.26 | 4.64 ± 0.25 | 1.42 ± 0.15 | 7.74 ± 0.30 | 0.72 ± 0.04 | 0.34 ± 0.00 |
| **bdh_a3** | 34.74 ± 0.50 | 4.89 ± 0.10 | 1.22 ± 0.08 | 7.81 ± 0.08 | 0.67 ± 0.07 | 0.35 ± 0.02 |
